# Incomplete information about the partner affects the development of collaborative strategies in joint action

**DOI:** 10.1101/374769

**Authors:** Vinil T. Chackochan, Vittorio Sanguineti

## Abstract

Physical interaction with a partner plays an essential role in our life experience and is the basis of many daily activities. When two physically coupled humans have different and partly conflicting goals, they face the challenge of negotiating some type of collaboration. This requires that both subjects understand their opponent’s state and current actions. But, how would the collaboration be affected if information about their opponent were unreliable or incomplete? Here we show that incomplete information about the partner affects not only the speed at which a collaborative strategy is achieved (less information, slower learning), but also the modality of the collaboration. In particular, incomplete or unreliable information leads to an interaction strategy characterized by alternating leader-follower roles. In contrast, more reliable information leads to a more synchronous behavior, in which no specific roles can be identified. Simulations based on a combination of game theory and Bayesian estimation suggested that synchronous behaviors denote optimal interaction (Nash equilibrium). Roles emerge as sub-optimal forms of interaction, which minimize the need to know about the partner. These findings suggest that physical interaction strategies are shaped by the trade-off of between the task requirements and the uncertainty of the information available about the opponent.

**Author summary:** Many activities in daily life involve physical interaction with a partner or opponent. In many situations they have conflicting goals. Therefore, they need to negotiate some form of collaboration. Although very common, these situations have rarely been studied empirically. In this study, we specifically address what is a ‘optimal’ collaboration and how it can be achieved. We also address how developing a collaboration is affected by uncertainty about the partner. Through a combination of empirical studies and computer simulations based on game theory, we show that subject pairs (dyads) are capable of developing stable collaborations, but the learned collaboration strategy depends on the reliability of the information about the partner. High-information dyads converge to the optimal strategies in game-theoretic sense. Low-information dyads converge to strategies that minimize the need to know about the partner. These findings are consistent with a game theoretic learning model which relies on estimates of partner actions, but not partner goals. This similarity sheds some light on the minimal computational machinery which is necessary to an intelligent agent in order to develop stable physical collaborations.

## Introduction

Many activities in daily life involve coordinating our movements with those of a partner or opponent. A couple of dancers, a couple of fighters, a team of players, two workers carrying a load or a therapist interacting with a patient are just the first examples which come to mind. In all these situations, each participant in the interaction needs to know what his/her partner is doing and/or intends to do. On this basis, he/she must then select their own action [1, 2]. To do this, the two partners (a ‘dyad’) may communicate verbally or non-verbally, or may watch each other. If the dyad participants are in physical contact, the forces they exchange are a rich source of information on their partner’s ongoing actions [3]. However, mechanical coupling also places restrictions on individual movements, so that the choice of what to do or not do must account for what the partner is doing. Specifically, stronger coupling is more informative but makes coordination more difficult, whereas weaker coupling facilitates coordination but provides less reliable information about partner’s actions [4]. These situations have been often studied in contexts in which there is one common and shared goal –for instance, control of isometric force [5, 6], reaching the same fixed [7, 8] or moving target [4, 9, 10], or operating a tool [3]. In these situations, dyads generally exhibit a better performance [7, 9] than a person performing the same task alone. Further, the improved performance resulting from training as a dyad also transfers to subsequent individual performance [9]. The advantage of dyad with respect to solo performance may be due to subdivision of efforts – less effort leads to less motor errors [11] – and the greater accuracy of a shared estimation of external events [9]. Specialized behaviors (i.e., ‘roles’) have been consistently observed within a dyad [12]. A common distinction is between leader and follower roles – leaders initiate the action and contribute most effort [6] – which are commonly observed in forms of interaction in which the coupling is acoustic or visual, like joint tapping [13] and mirror games [14] but have been also reported when the coupling is physical [15].

What happens if the information about our partner is incomplete or partial? For instance, when the two partners can only partly see or hear each other, or when they wear padded gloves which limit tactile perception? In this case we can expect a dramatic degradation in their ability to predict their partner’s actions. However, how does such uncertainty affect the decision of which action to perform? We can expect little differences when there is a shared goal – the action to perform is the same anyway. However, if the two partners have different and partly conflicting goals, and therefore they must negotiate a collaboration, how would uncertainty affect the emerging joint coordination? Although they are especially critical in social situations [2], these forms of interaction have received much less attention. These situations are characterized by a continuous interplay of two processes: understanding the ‘world’ – dyad and environmental dynamics, partner actions and possibly partner goals; and negotiating a mutually satisfactory coordination strategy. A series of studies [16, 17] focused on ‘motor’ versions of classic non-cooperative games (like the prisoner’s dilemma) – in which position-dependent force fields encoded the player-specific costs or rewards of the interaction – or very simple motor games (e.g. rope pulling). Bimanual versions of these tasks – in which there is only one controller – ended up in a cooperative solution. The dyad versions - two independent controllers – converged to the optimal non-cooperative solution – Nash equilibrium – a situation in which no partner can improve his/her strategy by acting unilaterally [18]. Bayesian estimation and game theory are the natural frameworks to address these scenarios, but have never been applied to joint action involving continuous coordination with physical coupling and conflicting goals.

Collaborative behavior, if any, is the end result of learning and adaptation through repeated performance, during which the agents gradually gain knowledge about dyad dynamics, the task requirements, and the partner’s actions. Nash equilibria describe optimal collaborative behaviors, but do not explain how they are achieved. Several mechanisms have been proposed [19] to account for learning a collaboration. They differ in terms of the assumptions on the partner’s actions and intentions are represented; for instance, at each trial each player may form beliefs about the opponent’s play and behaves rationally with regard to these beliefs – fictitious play [20, 21]. In this case, players don’t need to know about their opponent’s goal (i.e. their intentions); they just need to form beliefs about how their opponents will play (i.e. their actions). Alternatively, each agent forms a model of the opponent’s goals. The mechanisms through which collaborations are developed are as yet unclear and relatively unexplored.

In principle, learning to collaborate requires that both subjects know everything about their own and possibly their partner’s goals. In individual subjects, adaptation to a novel dynamic environment, for instance a new tool, requires reliable information on the consequences of own actions; incomplete information may slow down learning and/or may affect its outcome [22–25]. In dyads, if the information about the partner is partial or incomplete, optimal collaboration may be difficult to achieve [26]. It is unclear how partial information affects establishing a collaboration in two physically interacting humans.

Here we address how ‘optimal’ collaboration can be defined when two partners have partly conflicting goals. Further, we investigate how such collaboration can be ‘learned’. Finally, we address how the learned collaboration is affected by amount and quality of information about the partner.

## Materials and methods

### Experimental apparatus and task

Each experiment involved one pair of subjects (a dyad). Participants sat in front of two separate computer screens and grasped the handle of a three-dimensional haptic interface (Novint Falcon). They could not see or hear each other, and were not allowed to talk. The experimental apparatus is depicted in Fig. 1. The subjects were instructed to perform reaching movements in the vertical plane, between the same start point (displayed as a white circle, ⊘ 1 cm) and the same target point (yellow circle, ⊘ 1 cm), but through different via-points. In a reference frame centered on the robot workspace (one for each subject), with the X axis aligned with the left-right direction and the Y axis aligned with the vertical direction, the start point was placed in the (−5, 0, 0) cm position and the target point was placed in the (5, 0, 0) cm position. Hence the start and the target point had a horizontal distance of 10 cm. The subjects were also instructed to keep their movements as planar as possible, i.e. by keeping the depth, Z coordinate within the 18-26 cm range with respect to the origin of the workspace. The current positions of the end effectors, *x*_1_ and *x*_2_, were continuously displayed to each partner, as 0.5 cm circular cursors on their respective screens, colored in green if the depth was correct and in red otherwise. A trial started when both subjects placed their cursor inside the start region. Then the target and a via-point (⊘ 0.5 cm circle) appeared. The via-points were different for the two subjects and were placed, respectively, at locations VP_1_ = (−3,−2,0) cm and VP_2_ = (3,2,0) cm. The haptic interfaces generated a force proportional to the difference of the two hand positions:

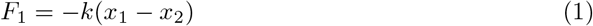

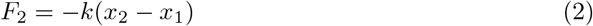

with *k* = 150 N/m. Hence, the two subjects were mechanically connected. At the end of each movement, each subject received a 0-100 reward, calculated as a function of the minimum distance of their movement path from his/her own via-point and of the average interaction force:

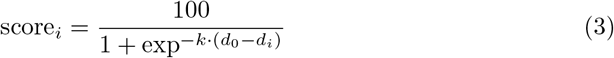

where *d_i_* = *d_V Pi_* + c · d_12_ and *i* = 1, 2. The quantities *d_V Pi_* and *d*_12_ are, respectively, the minimum distance between the movement trajectory and the subject’s own ‘via-point’ (VP_*i*_) and the average distance between the two subjects’ hand positions. In the disconnected trials we took *c* = 0, i.e. the score only depended on how close the subjects got to their own via-point. Parameters *k* and *d*_0_ were calculated so that the score was maximum (100) for *d_i_* ≤ 0.005 m (i.e., the VP radius), and minimum (0) for d_i_ ≥ 0.02 m. Audio cues were provided at the start and end of the movements. To encourage subjects to establish a collaboration, in trials in which the two subjects were mechanically connected we took *c* = 0.5, so that in order to get a maximum score subjects also had to keep their relative distance as low as possible. The subjects were instructed to aim at maximizing this score. Specifically, they were told that performance depends on how close they would get to the via-point. They were also told that they might experience a force while performing the task. They were warned that force magnitude also affects the score, and were instructed to also keep this force to a minimum. To encourage subjects to maintain an approximately constant movement duration, after each movement a text message on the screen and changes in the color of the target (either green or red) warned the subjects if the movement was either too fast (duration < 1.85 s) or too slow (duration > 2.15 s). However, the participants received no penalization if their movement duration did not remain within the recommended range. The subjects pairs were randomly assigned to three groups depending on the feedback provided about the interaction force. In the haptic (H) group, interaction could only be sensed haptically. In the visuo-haptic group (VH), interaction force (magnitude, direction) was also displayed as an arrow attached to the cursor (scale factor: 10 N/cm). In the partner-visible group (PV), subjects could see their partner’s cursor. Therefore, these subjects have a more reliable information about partner’s movement. The experiment was organized into epochs of 12 movements each. The experimental protocol consisted of three phases: (i) baseline (one epoch), (ii) training (ten epochs) and (iii) after-effect (two epochs) for a total of 13 × 12 = 156 movements. During the baseline phase the interaction forces were turned off, and each subject performed on their own. During the training phase the subjects were mechanically connected. During this phase, in randomly selected trials (catch trials) within each epoch (1/6 of the total, i.e. 2 trials per epoch) the connection was removed. The connection was permanently removed during the after-effect phase. During the training phase the subjects had the option to establish a collaboration - negotiating a path through both via-points, which would lead to a minimization of the interaction forces and a maximum score for both - or to ignore each other - each partner would only focus on their own via-point and on maximizing his/her own score. We developed a custom software application using CHAI3D, an open source framework for control of haptic devices [27].

**Fig 1.**
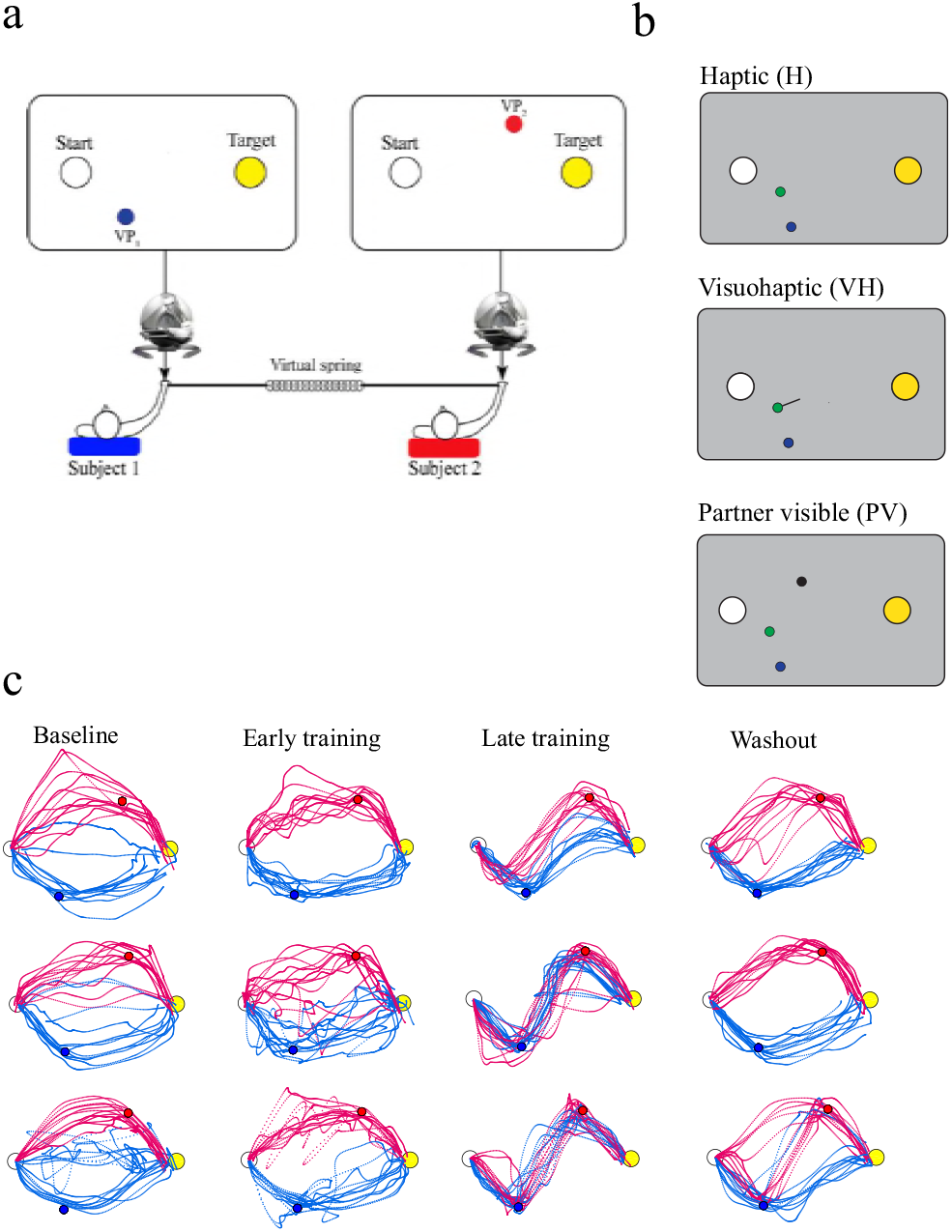
Experimental apparatus and protocol. **(a)**, Partners in a dyad were connected through a virtual spring. Both subjects were instructed to perform reaching movements in the vertical plane, between the same start point and the same target point but through different via-points (VP). Each subject could only see his/her own VP, but not their partner’s. Both were instructed to keep the interaction force as low as possible during movement. The experimental protocol consisted of three phases: baseline, training and after-effect. During the baseline phase the interaction forces were turned off, and each subject performed on their own (‘solo’ performance). The subjects were mechanically connected during the training phase, and the connection was permanently removed during the after-effect phase. **(b)** We manipulated the information available on partner’s actions by providing it either haptically, through the interaction force (Haptic group, H) or by additionally displaying the interaction force vector on the screen (Visuo-Haptic group, VH) or displaying partner’s cursor itself (Partner Visible group, PV). The yellow and white circles denote, respectively, the start and target position. The green circle is the cursor location. In the VH group, direction and magnitude of the interaction force is depicted by a line originating from the cursor. In the PV group, partner’s cursor is shown by black circle. **(c)**. Movement paths in baseline (unconnected), early-training, late-training and washout phases of the experiment from three typical H, VH and PV group dyads

**Fig 2.**
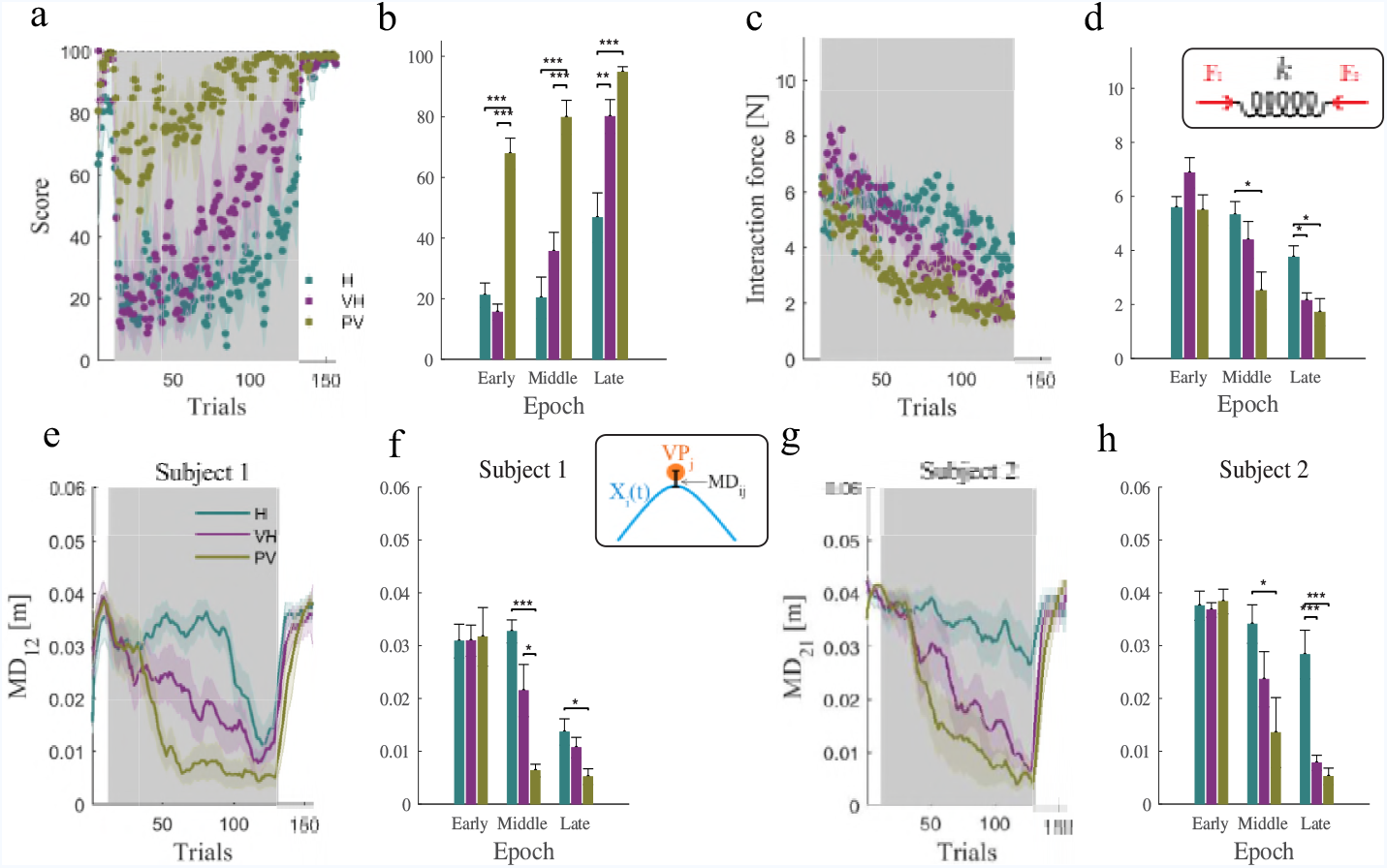
Dyad develop a collaboration, and learning depends on the amount of information available about the partner. **(a)**, Temporal evolution of score over trials, for the haptic (H), visuo-haptic (VH) and Partner-visible (PV) groups respectively. **(b)**, Score at the beginning, middle and at the end of training. **(c)**, Magnitude of the interaction force over trials. **(d)**, Interaction force at the beginning, middle and at the end of training. **(e)**, Magnitude of the distance from partner VP for subject 1 over trial for the H, VH and PV groups. **(f)**, Minimum distances of subject 1 from his partner VP at the beginning, middle and at the end of training. **(g,h)**, As in **(e,f)** for subject 2. The areas in grey denote the training phase. All plots indicate the population means. Error bars and shaded areas denote the standard error (SE). Asterisks indicate statistically significant differences (\**P <* 0.05, * \**P <* 0.01, * * * *P <* 0.001)

### Subjects

A total of 30 subjects participated in this study, recruited among the graduate and undergraduate students of University of Genoa. All subjects were right-handed, as assessed using the Edinburgh Handedness Inventory [28], naΪve to the task and with no known neurological or motor impairment at the upper limb. From the list of participants, we formed 15 dyads with similar body size (assessed through the body mass index) which were randomly assigned to the H (25 *±* 5 y; 9 M + 1 F), VH (24 *±* 3 y; 8 M + 2 F) and PV groups (24 *±* 3 y; 6 M + 4 F). The research conforms to the ethical standards laid down in the 1964 Declaration of Helsinki that protects research subjects and was approved by the competent ethical committee (Comitato Etico Regione Liguria). Each subject signed a consent form conforming to these guidelines.

### Data Analysis

Hand trajectories and robot-generated forces were sampled at 100 Hz and stored for subsequent analysis. The data samples were smoothed by means of a 4th order Savitzky-Golay filter with a 370 ms time window. We used the same filter to estimate velocity and acceleration. We identified the start and end times of each trajectory as, the time instants at which the speed crossed a threshold of 2 cm/s. In the analysis, we specifically focused on the temporal evolution of the trajectories and on signs of collaboration between partners within the same dyad. Collaboration can be characterized in terms of both movement kinematics and movement kinetics. Interaction force (IF) is calculated as 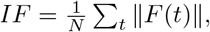 where *F*(*t*) is equal and opposite for the two partners in the dyad – see Eq. 2. Less interaction force would point at a greater collaboration.

A sign of collaboration is that each subject, while passing through his/her own via-point, also gets very close to his/her partner’s. This can be quantified in terms of the Minimum via-point distance (MD_*ij*_), defined as the minimum value of the distance of subject *i* to the j-th via-point: *MD_ij_* = min_*t*_||*x_i_*(*t*) − *x*VP_*j*_|| with *i, j =* 1, 2. If *i* ≠ *j* this quantity reflects how close each subject gets from his/her partner’s via-point.

Looking at the power developed by each subject would provide information on whether the subjects move actively, or are passively pulled by their partner through the mechanical coupling. To quantify this, we calculated the power (P_*i*_), defined as the scalar product of the interaction force *F_i_*(*t*) and the velocity vector *v_i_*(*t*) of each of the subjects. At a given time, a negative power would mean that the subject is controlling his/her motion (i.e. he/she is behaving as a ‘leader’). Conversely, a positive power would indicate that the subject is being pulled toward the other (i.e., he/she is behaving as a ‘follower’) – see Fig. 3(c). We specifically focused on the average power calculated in the 300-ms interval taken just before and just after the crossing of each via-point. We denote as LI_*ij*_this value for the *i*-th subject and the *j*-th via-point.

**Fig 3.**
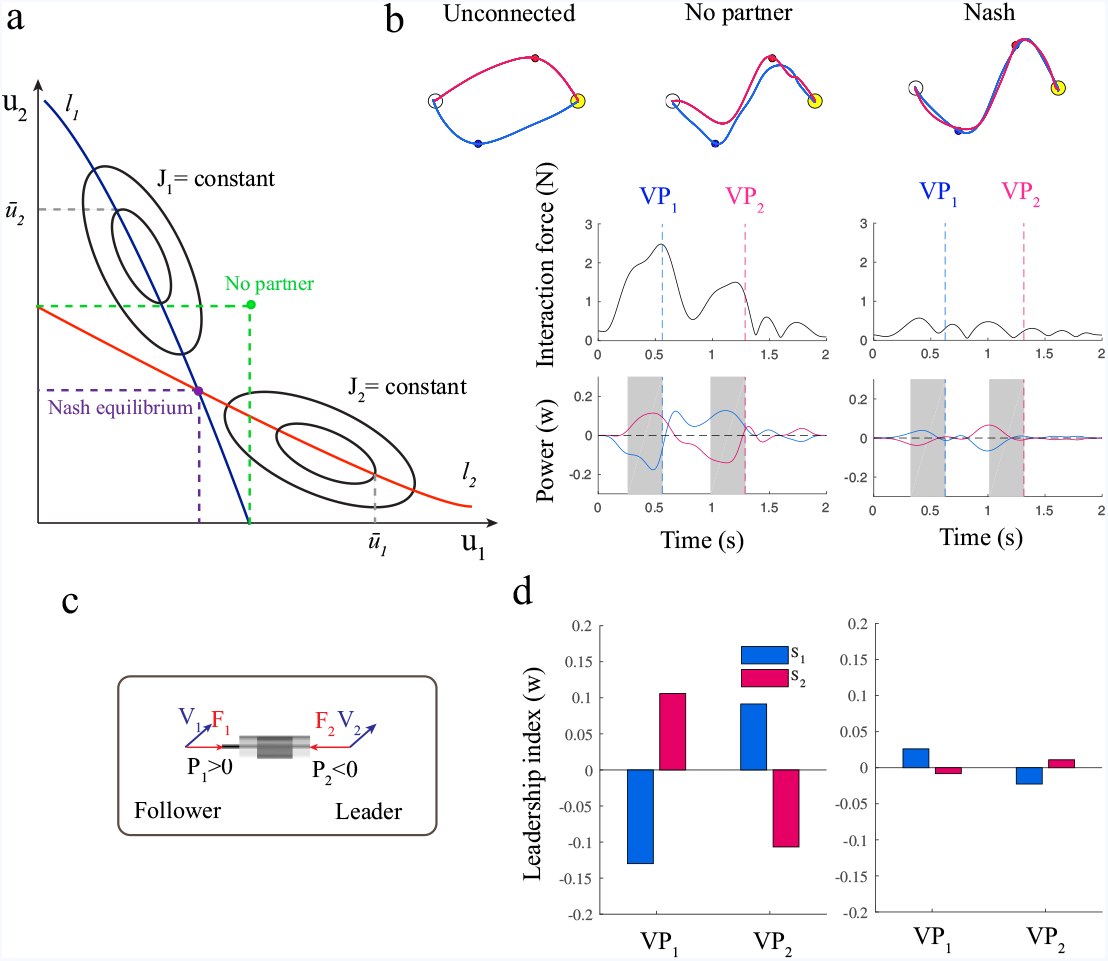
Predictions based on game theory. **(a)**, Definition of Nash equilibria vs ‘no-partner’ strategies in two-agents non-cooperative game. Nash equilibrium is determined by the intersection of the partners’ reaction curves - locus of optimal control action calculated for each value of the partner action [29] (blue and red lines). The ‘no-partner’ solution is determined as the optimal action calculated by each agent by assuming that partner’s control is zero. **(b)**, Simulated movements under Nash equilibria vs. ‘no-partner’ conditions, for subject 1 (blue) and subject 2 (red). From top to bottom: movement paths, interaction force and interaction power profiles. **(c)** Interaction power, *P_i_*, is defined as the scalar product of the interaction force (*F_i_*) and the velocity(*V_i_*). **(d)**, Leader-follower strategy in the no-partner (left) and Nash strategy (right). The plots depict the Leadership index (LI) for both subjects and both via-points, calculated as the average power in the 300 ms interval (in gray) just before crossing the via-point by its homologous subject

We expect that task performance at subjects and dyad level evolves with time (learning) and is affected by the amount of information each subject has available about his/her own partner. To test this, for all the above indicators we ran a repeated-measures ANOVA with group (H, VH, PV) and Time (early - epoch 1, middle - epoch 6 and late - epoch 11) as factors. To further analyze the group differences (if any), we then compared (planned comparisons) the between-groups differences (H and VH) for each Time value (early, middle, late). All data were analyzed using MATLAB and the statistical tests were performed using R. Statistical significance was considered at *P <* 0.05 level for all tests.

### Computational Model

We compared the observed movements with the outcome of a computational model, whose purpose was to predict the ‘optimal’ behaviors and how they depend on the information available about the partner. We approximated the dyad dynamics and the subjects’ sensory systems as a linear discrete-time dynamical system with additive Gaussian noise in both motor commands and sensory measurements. In particular, dyad dynamics was approximated as a pair of point masses connected by a spring. We assumed that each subject operates his/her own point mass by applying a force to it. We also assumed that each partner’s sensory system provides visual or proprioceptive information about his/her own position, plus haptic information about the interaction force. The interaction strategy is completely specified by a pair of feedback controllers (one per partner). The task was specified by a pair of quadratic cost functions (one per partner); see the Supplementary Note for details. The possible strategies can be studied in terms of non-cooperative game theory – where non-cooperation means that the two partners develop their strategies independently. The optimal non-cooperative solution – Nash equilibrium [18] – corresponds to a situation in which each partner cannot improve his/her strategy unilaterally; see Fig. 3(b). Nash equilibrium strategies of feedback type can be computed in terms of differential game theory [29] in the same way optimal feedback control theory has been used to predict optimal movements of a single human [30]. Nash equilibria represent the optimal form of collaboration which two partners can achieve when independently planning their actions (i.e. non-cooperatively). Importantly, these strategies require that each agent has a perfect knowledge of his/her partner’s cost function. Another possibility is that the two partners do not collaborate at all, in the sense that they ignore each other when determining their control policy. We modelled this situation (no-partner) by assuming that each subject develops a control strategy by considering the partner’s control as noise. In this case, the problem reduces to separately developing two independent optimal LQG controllers. In this case, each agent only needs to know his/her own cost function. The control strategies are not the only determinants of behavior. Both partners have incomplete knowledge of the system state, therefore they need to estimate it by means of a ‘state observer’ [31]. A state observer relies on the optimal combination of prediction and correction. Prediction requires an accurate model of dynamics and a copy (efferents copy) of his/her own motor command. Correction is driven by the information provided by the sensory system. However, in the case of a dyad, an accurate state prediction is only possible if each agent has information on the partner’s motor command. In other words, each agent must know what the other partner is doing.

Using a computational model involving separate controllers and state observers – see Fig. 5(a) – we simulated the iterative learning process based on fictitious play for all three experimental conditions (H, VH and PV). For each trial, each subject developed a model of their partner’s control action, which was incorporated in their updated control policy – see the Supplementary Note for details. The simulation results were analyzed in exactly the same way as the experimental results. In particular, for both models we calculated both dyad- and subject-level indicators. All simulations were performed in MATLAB/Simulink.

## Results

We designed a novel interactive task in which – somewhat resembling the classic ‘battle of sexes’ game [32] – players must reconcile different goals with a common preference to stay close together. Two subjects were mechanically connected but could not see each other. They were instructed to perform reaching movements with the same start and end positions, but through different via-points (VP); see Fig. 1a. Both subjects were also instructed to keep the interaction force as low as possible during movement. They had the option of establishing a collaboration - negotiating a path through both VPs, which would lead to a minimization of the interaction forces - or to ignore each other, by only focusing on their own goal. We manipulated the information available on partner’s actions by providing it either haptically, through the interaction force (haptic group, H); by additionally displaying the interaction force vector on the screen (visuo-haptic group, VH); or by continuously showing the partner movements (partner visible, PV); see Fig. 1b.

## Collaboration in dyads and the role of information

In all three haptic (H), visuo-haptic (VH) and partner-visible (PV) groups, all dyads converged to stable and consistent behaviors; see Fig. 1c-e. When the connection was removed, both agents quickly returned to the baseline situation. These observations are confirmed when looking at the score, the interaction force and the minimum distance from the partner’s VP. All quantities are expected to improve if subjects establish a collaboration.

The temporal evolution of score for subject pairs is summarized in Fig. 2a. Subjects in the VH group achieved a greater score than those in the H group at the end of training, which is confirmed by statistical analysis. Overall the subject pairs improved their movement score with training (*F*_2,24_ = 47; *P <* 10^*−*4^) and exhibited significant group differences (*F*_2,12_ = 56.07; *P <* 10^*−*4^). We also found a significant group*×*time interaction (*F*_4,24_ = 5.2; *P* = 0.0034). Post-hoc analysis confirmed that subject pairs in the VH group achieved a significantly greater score at the end of training phase than the H group (*P* = 0.004). Also the PV group differs significantly from H (*P <* 0.0002). At middle time, PV again differs not only from H (*P <* 10^*−*3^), but also from VH (*P <* 10^*−*3^), see Fig. 2b.

The interaction force is a major determinant of the score, and its temporal evolution exhibits a similar behaviour in the three groups – see Fig. 2c. Overall, we found a significant training (time) effect (*F*_2,24_ = 37.4; *P <* 10^*−*4^), a significant group effect (*F*_2,12_ = 6.2; *P* = 0.014), and a significant group × time interaction effect (*F*_2,12_ = 6.2; *P* = 0.014) – see Fig. 2d. Post-hoc analysis showed that groups VH-H (*P* = 0.04) and PV-H (*P* = 0.01), but not PV-VH, differ significantly at late time. In addition, groups PV-H (P=0.02) but not VH-H and PV-VH differ significantly at middle time. In summary, the trial-by-trial improvement of both score and interaction force is faster in the PV group and slower in the H group.

A similar behavior can be observed in the temporal evolution of the minimum distance from the partner’s via-point is depicted in Fig. 2e-h. In both groups and in both subjects in the dyad, the minimum distance (MD) decreases over trials and quickly washes out when the connection is permanently removed (after-effect phase). The magnitude of the decrease is very similar in all groups. Statistical analysis confirmed this observation. We found a significant time effect for both subject 1 (*F*_2,24_ = 40; *P <* 10^*−*4^) and subject 2 (*F*_2,24_ = 36.4; *P* = 0.0067). We also found significant group effects (*F*_2,12_ = 40; *P <* 10^*−*4^) for subject 1 and (*F*_2,12_ = 8.9; *P* = 0.004) for subject 2. However, we only found a significant group×time interaction for subject 2 (*F*_2,16_ = 5.64; *P* = 0.014), but not for subject 1. Post-hoc analysis showed that for subject 1, in the groups (PV-H) the MD value is significantly different (lower in the PV group) in the late time (*P* = 0.0296) and also group combinations (PV-H and PV-VH) differ significantly at the middle time (*P* = 0.0002, *P* = 0.01 respectively). For subject 2, post-hoc analysis showed that group pairs (VH-H and PV-H, but not PV-VH) differ significantly at the late time (*P* = 0.0009, *P* = 0.0003 respectively) and groups (PV-H, but not VH-H and PV-VH) significantly differ at the middle time (*P* = 0.04). In other words the three groups – specially H and PV – differed in both magnitude and rate of decrease of their via-point distance. Fig. 2f, h, summarizes the effect of learning in terms of MD in three groups. This also suggests that in the H group learning is less complete for subject 2 than subject 1. Overall, these results suggest that in the PV group learning is faster and results in a better performance, followed by VH and then H (greater score, lower interaction force, lower distance from partner VP).

## Optimal interaction and the emergence of roles

The above results still say little on the nature of the collaboration and how the collaboration is developed. To address this, we developed a computational model, based on differential game theory [29], to predict the ‘optimal’ interaction behaviors – see the Methods section for details. Consistent with computational models of individual movements base on optimal control [30], the interaction strategy is completely specified by a pair of feedback controllers (one per partner). Using the model, we simulated an optimal collaboration (Nash equilibrium) [18], in which no partner can improve his/her strategy unilaterally. Another possibility is that the subjects determine their control actions by assuming that they are alone in controlling the dyad dynamics. As a consequence, they focus on their own via-point and on minimizing the interaction. In this case, information on what the other partner is doing is not accounted for during action selection. This alternative scenario defines the maximum compliance with the task achievable with the minimum amount of collaboration between partners. We refer to this scenario as the ‘no-partner’ strategy – see Fig. 3a.

Fig. 3b summarizes the model predictions. The movement trajectories look similar in the two models, but a closer look suggests that in the no-partner case each subject actively moves toward his/her own via-point – thus behaving as a ‘leader’, but is pulled by the partner when getting closer to the other via-point – thus switching to a ‘follower’ role. This effect is clearly visible when looking at the average interaction power calculated just before crossing the via-point – see Fig. 3. As a consequence, the no-partner scenario exhibits temporal delays between the via-points crossing times and a greater magnitude of interaction force and interaction power. In contrast, in the optimal (Nash) scenario the two subjects approximately follow the same trajectory, by crossing each via-point at approximately the same time. Both the interaction force and the interaction power remain low over the whole movement, and there are no clear leader-follower roles. Therefore, a distinctive feature of the ‘no-partner’ scenario is the alternation of leader and follower roles - each subject acts as a leader when crossing his/her own via-point, and as a follower when crossing that of the partner. This is also reflected in the different crossing times (with respect to the leader, the follower lags behind). In conclusion, establishing roles can be seen as a form of compensation for poor integration of the partner’s intentions into the subject’s own control strategy.

Based on these predictions, we looked into the emergence of distinct leader-follower roles in our experimental data at the end of the training phase. Fig. 4a,b summarizes the leadership indices (*LI*) – average interaction power in the 300 ms interval before via-point crossing – calculated in the late epochs at *V P*_1_ (a) and *V P*_2_ (b). As regards *LI*_12_ *− LI*_22_ – difference in leadership indices for subject 1 and subject 2 at *V P*_2_ – we found significant group differences (*F*_2,12_ = 5.84; *P* = 0.016). Planned comparison confirmed significant differences between H and VH (*t*_7.75_ = 2.63*, P* = 0.03), H and PV (*t*_7.31_ = 3.29*, P* = 0.01) but not VH and PV (*P* = 0.43). Also, we found no significant effects for difference in leadership indices for subject 1 and subject 2 at *V P*_1_. These results indicate that when there is limited information about the partner (group H), Subject 1 exhibits a transition from a leader role near *V P*_1_ to a follower role near *V P*_2_. The effect decreases and tend to vanish when the amount of available information about the partner increases (from H – minimum information – to PV – maximum information). Although not statistically significant, Subject 2 exhibits a similar trend – leader near *V P*_2_ and follower near *V P*_1_. When comparing these results with the simulations, at the end of the training phase the dyads in the PV group are more similar to the optimal (Nash) strategy. To compare model predictions and experimental results, we calculated the difference in the interaction power for both partners at *V P*_1_ and *V P*_2_ and for the corresponding Nash (green) and No-partner (yellow) scenarios – see Fig. 4b. Overall, the experimental results suggest that dyads with more available information (PV group) about the partner are closest to the optimum (Nash) scenario, whereas dyads with less reliable information (H group) resemble more the no-partner scenario.

**Fig 4.**
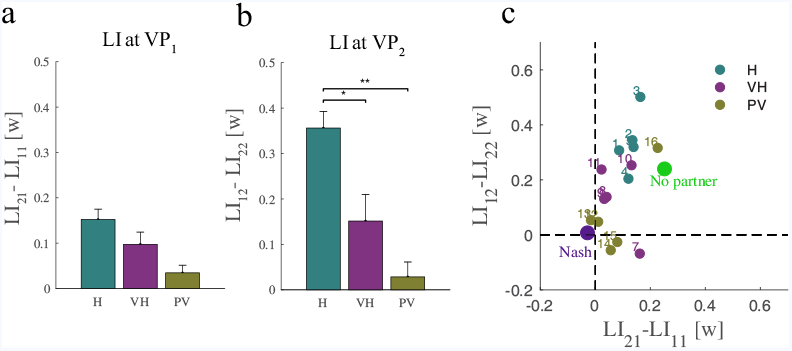
Leadership Index (LI) differences at the end of training (average within the last training epoch). **(a)**, Δ*LI* at *V P*_1_ is calculated as the difference between *LI*_21_ and *LI*_11_. **(b)** Δ*LI* at *V P*_2_ is calculated as the difference between *LI*_12_ and *LI*_22_. **(c)**, Δ*LI* for each dyad in all groups. The large dots denote the No partner (green) and the Nash (violet) model predictions. The plots indicate the population means. Error bars denote the standard error (SE). Asterisks indicate statistically significant differences (*\*P <* 0.05, *\* * P <* 0.01, *\* * * P <* 0.001)

## Learning to collaborate

Consistent with computational models of sensorimotor control of individual movements [33], we posited that each partner uses a state observer to predict the dyad state from sensory and motor information. The state observer is easily extended to also account for the partner’s control action; see Fig. 5a. We simulated the process of establishing a collaboration through repeated performance. The model uses a form of ‘fictitious play’ [20, 21], which requires minimal assumptions on each player’s internal representation of their opponent [19]. At every trial, each subject independently estimates the most likely partner’s action and incorporates it into his/her own control policy on the next trial. This model is attractive as it requires minimum information about the partner – it does not need to establish a model of the partner’s task or goals. We simulated all three scenarios (H, VH, and PV) and found – see Fig. 5 – that more information leads to a more Nash-like collaboration, characterized by greater synchronization and less distinct roles. This is confirmed when looking at the leadership index; see Fig. 5. Switching of roles (each subject leads when aiming at his/her own VP and follows when aiming at the partner VP) – which denotes a lack of consideration of partner intentions when developing their own control policy – decreases as the amount of information about the partner increases.

**Fig 5.**
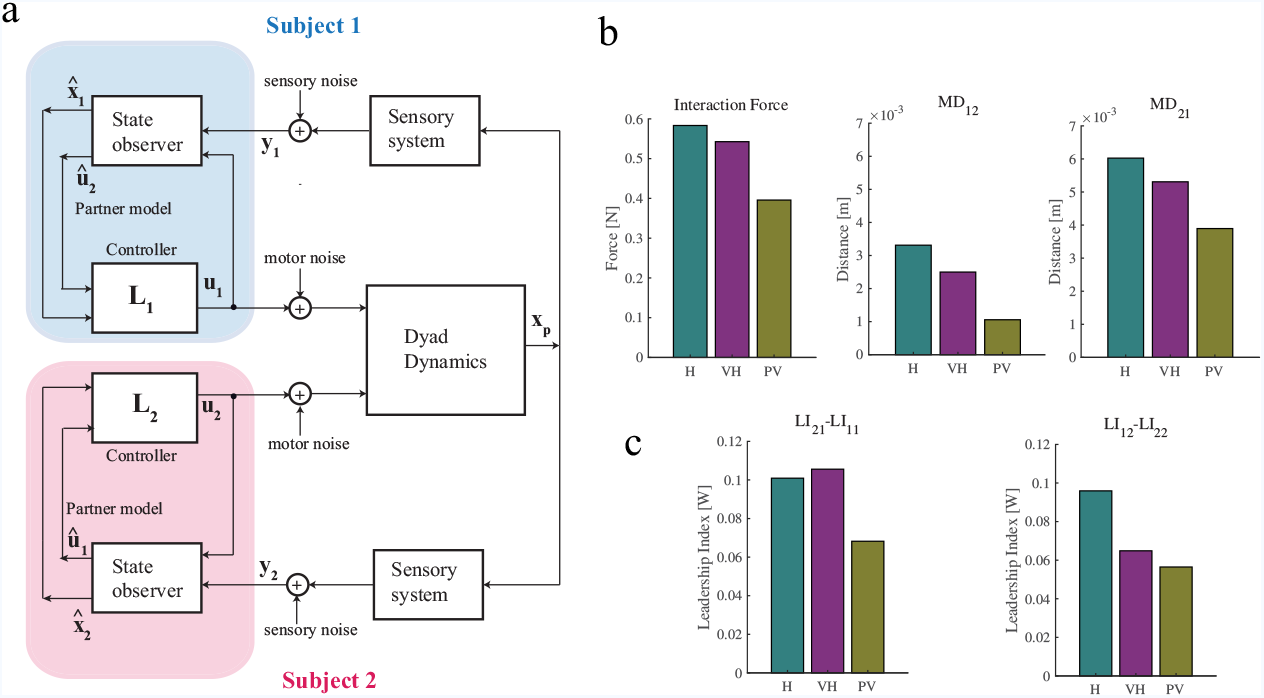
**(a)**, Computational model. Two partners jointly control dyad dynamics (approximated as two mass-points connected by a spring). Each partner has a separate sensory system which provides information about dyad state. The controller includes a state observer, a feedback controller and an estimate of the partner controller (partner model). **(b)**, From left to right: Interaction force and minimum distance from *V P*_1_ (*MD*_21_) and *V P*_2_ (*MD*_12_), for all three H, VH and PV scenarios. **(c)**, Leadership index for subject 1 and subject 2 at *V P*_1_ (left) and *V P*_2_ (right) for all three H, VH and PV scenarios

## Discussion

In individual sensorimotor control, uncertainty affects the estimation of the state of the body and the external environment (including tools, if any). Inaccurate state estimates may lead to inaccurate or inefficient control. In joint action, uncertainty may also affect estimation of the opponent’s ongoing actions (and possibly their ultimate goals). This may make establishing a collaboration more difficult or even impossible. Physical coupling is a major source of information about the opponent’s behaviors. While operating a pole by pulling ropes [3], dyads produce more overlapping forces than individuals using both hands. In dyad performance, subjects must keep their rope stretched to collect information about their partner’s action. If they don’t – for instance, when the rope is loose – they just have no way to coordinate their movements. These results suggest that subjects need information about their partner in order to establish a collaboration. Stronger coupling is more informative but makes coordination more difficult, whereas weaker coupling facilitates coordination but is a less reliable source of information [4].

We argued that the development of an effective interaction is profoundly affected by information uncertainty. To test this, we focused on three groups of dyads, characterized by different amounts of information about their opponent. The Haptic (H) group only relied on haptic information – from a relatively weak coupling, hence a rather unreliable information. The other groups (Visuo-Haptic, VH and Partner-Visible, PV) were provided with increasingly rich and reliable information about their opponent. We also developed a computational model of the interaction, in which the mechanically coupled players form a single mechanical system, which they must jointly control. Consistent with the optimal feedback control [30] and optimal Bayesian estimation frameworks [31], which are widely used in modeling single-player sensorimotor control, we assumed that each subject has his/her own optimal feedback controller, completely specified by his/her own assigned task, and a state (and partner) observer, which combines sensory information and predicted dyad dynamics to estimate the dyad’s internal state and the partner actions – see Fig. 3a. The model summarizes the available knowledge on the neural basis of joint coordination [33], but has been applied to the study of joint interaction in very limited situations, involving either discrete decisions [17] or shared (e.g. bimanual) control [34]. The application of this modeling framework to joint sensorimotor control is one major novelty of the present study.

In our proposed control model we show that reliable estimation of partner’s action can be achieved through a simple extension of the state observer. In other words, we assume that partner’s action recognition is obtained through a optimal (in Bayes’ sense) combination of predictions and observations. Our results provide no information on the possible neural substrates of action observation, which is typically associated with the mirror neuron system [35]. However, our prediction that estimation of partner actions during joint action is no different from estimating other aspects of plant state points at a role of the cerebellum (which is involved in the representation of body and environment dynamics), in conjunction with brain areas like the superior temporal sulcus that have been associated to action observation [2].

### Dyads gradually develop stable strategies

Our results suggest that dyads in all three groups gradually converge to stable interaction strategies, characterized by low interaction forces and low distances from the partner’s via-points. When two subjects are rigidly coupled, the development of stable coordination is relatively fast [12] or even instantaneous [8]. When the coupling is softer, learning is more gradual [4, 9]. The rapid emergence of stable coordination in rigid coupling may be due to the fact that the partner actions are easier to predict and a joint coordination strategy is simple to develop as the two subjects shared the same task. However, dyads need more time to develop an interaction strategy when the task is more challenging [4, 10] or – as in the present study – when the subjects have different and partly conflicting goals.

### Differential game theory as general model of joint coordination

The application of game theory concepts is relatively novel in the study of joint sensorimotor interaction. Few studies have focused on motor versions of classical games, like the prisoner’s dilemma and rope pulling [16, 17]. Although these tasks involved movements, their primary focus was on discrete decisions. These games have distinct cooperative and non-cooperative solutions, in which the players either agree or individually determine their actions. Game theoretic concepts are implied – although not explicitly stated – in the notion of effort sharing [36]. The present study focuses on a purely motor task and points at differential game theory as a general modeling framework for joint coordination. Hence our analysis can be easily extended to a broad range of tasks and situations. The present study specifically uses game theory to study optimal forms of interaction and to contrast them with possible alternatives. Our focus is exclusively on non-cooperative situations - in fact, in our task the cooperative and non-cooperative strategies are not distinguishable.

We used the computational model not only to predict optimal behaviors, but also to understand how these behaviors are learned. The study results suggest that fictitious play may account for the development of optimal collaboration. This model adds computational substance to previous models [37] of joint action, which only address the representation level. Our learning model specifically predicts that dyads converge to a Nash equilibrium if players have reliable information about their opponent. In contrast, if there is more uncertainty on the partner action, the dyad establishes a pragmatic form of collaboration, which requires minimal interaction with the partner. Our proposed model also has possible technological implications, as it sheds some light on the minimal computational machinery which is necessary to an intelligent agent in order to develop stable physical collaborations.

### The learned interaction is influenced by the amount of available information

Previous studies on joint coordination generally assumed that players had either full or complete lack of knowledge about opponent’s goals, state and ongoing actions, and did not explicitly address the dynamics of the learning process. The present study for the first time addresses the mechanisms underlying the development of collaboration when the information about the partner is incomplete.

We specifically observed that incomplete information prevents more efficient forms of collaboration, which require reliable estimates of the opponent’s state and current actions. The computational model allowed to identify one specific signature of the extent to which subjects use information about their partner’s actions when planning their movements. In simulations in which each subject computes his/her control action by ignoring the partner, subjects alternate leader and follower roles within the same movement. Subject 1 leads in via-point 1 and then follows his partner in via-point 2. Conversely, Subject 2 follows his partner in via point 1 and leads at via point 2. This behavior can be interpreted in terms of the ‘minimum intervention principle’ of optimal feedback control [30]. For Subject 1, via-point 2 is task-irrelevant and therefore getting close to this point is not controlled explicitly; vice versa for Subject 2.

These predictions are confirmed by our experimental results. We found that dyads characterized by less reliable partner information (group H) are qualitatively very similar to the predicted ‘no-partner’ behaviors. In contrast, leader-follower patterns disappear when information is more reliable. The dyads with more information about the partner (PV group) exhibit a form of collaboration which is very close to optimal (Nash strategy). Consistent with these observations, leader-follower roles are observed in imitation tasks, like joint tapping [13] or mirror games [14], when partners lack feedback about their opponent’s actions. Are roles affected by training? In a study involving joint generation of isometric forces [6], leader-follower relations were observed in novice-experienced pairs but were not affected by practice. In contrast, in both simulations and experiments, we found that roles evolve with the knowledge gained about the partner. As a consequence, in all groups, early trials exhibit distinct roles. In high-information dyads (VH and PV group) roles gradually disappear and the collaboration strategy comes close to Nash equilibrium. In contrast, in low-information dyads (H group) roles are preserved until the end of training. Overall, these observations suggest that leader-follower strategy is a sub-optimal form of collaboration, consequent to an incomplete co-activity.

### Do partners understand each other intentions?

The study results and the model simulations are consistent with the notion that each subject establishes a model of the opponent’s current actions. Similarly, in a sensorimotor coordination game, human players against computer opponents adapted their behavior to the opponent’s willingness to cooperate [38]. One crucial question in joint action is whether and to what extent the two partners within the dyad develop a deeper form of understanding, related to their partner’s goal. While there is some evidence [39] that subjects in a dyad do develop and maintain models of their opponent goals even when not strictly necessary (e.g. when acting individually), and even when this is detrimental to individual performance, other studies [40, 41] report little evidence of players modeling their partners’ goals or intentions.

In our study, the gradual decrease in the leadership indices suggests that subjects incorporate information about their partner into their motor plans. Our experiment was carefully designed so that subjects had no explicit clue on the partner’s task. After the end of each experiment, the participants gave no consistent answer when asked what they thought the partner was doing. This is consistent with the fictitious play model of learning, which does does not require to model the partner’s task explicitly – this is what should properly be referred as intention – but requires to account for the partner’s most likely action, inferred from previous trials. Therefore, both experiments and simulations suggest that at least in this task, subjects only need minimal information about their partner to converge to quasi-optimal behaviors (Nash equilibrium). However, these findings do not rule out the possibility that in other more complex forms of interaction the subjects estimate their partner goals. Indeed, the proposed Bayesian model of action estimation can be easily extended to estimation of partner’s goals. Future experiments, possibly involving generalization to other tasks or interacting with a virtual partner will be necessary to clarify this important point.

## Supporting information

**S1 File** Computational model of joint action coordination

**S1 Data** Zip file containing data from all indicators and all groups (H,VH, PV) - one file per indicator and per group. Each file has 5 rows (one per dyad) and 156 columns (one per trial).

## Acknowledgments

This research was supported by a grant from the Italian Ministry of University, Education of Research under the Research Projects of National Interest (PRIN) program (ModuLimb).

